# The Chordate Origins of Heart Regeneration

**DOI:** 10.1101/2023.09.19.558507

**Authors:** Keaton J. Schuster, Lionel Christiaen

## Abstract

The human heart is infamous for not healing after infarction in adults, prompting biomedical interest in species that can regenerate damaged hearts. In such animals as zebrafish and neonatal mice, cardiac repair relies on remaining heart tissue supporting cardiomyocyte proliferation. Natural *de novo* cardiogenesis in post-embryonic stages thus remains elusive. Here we show that the tunicate *Ciona*, an ascidian among the closest living relatives to the vertebrates, can survive complete chemogenetic ablation of the heart and loss of cardiac function, and recover both cardiac tissue and contractility. As in vertebrates, Ciona heart regeneration relies on Bone Morphogenetic Protein (BMP) signaling-dependent proliferation of cardiomyocytes, providing insights into the evolutionary origins of regenerative cardiogenesis in chordates. Remarkably, prospective lineage tracing by photoconversion of the fluorescent protein Kaede suggested that new cardiomyocytes can emerge from endodermal lineages in post-metamorphic animals, providing an unprecedented case of regenerative *de novo* cardiogenesis. Finally, while embryos cannot compensate for early losses of the cardiogenic lineage, forming heartless juveniles, developing animals gain their regenerative ability during metamorphosis, uncovering a fundamental transition between deterministic embryogenesis and regulative post-embryonic development.

## Introduction

In adult humans, damage to the heart such as myocardial infarction often results in chronically reduced cardiac function and poor prognosis due to the formation of fibrous non contractile scar tissue, instead of healing by replacing lost cardiomyocytes. These medical challenges have fostered interest in model organisms that present the natural ability to regenerate cardiomyocytes after injury. In vertebrates including zebrafish (Poss et al., 2002), neonatal mice (Porrello et al., 2011) and pigs (Ye et al., 2018), and amphibians (Becker et al., 1974; Laube et al., 2006; Marshall et al., 2019), the damaged heart can regenerate via dedifferentiation and proliferation of remaining cardiomyocytes (Jopling et al., 2010; Kikuchi et al., 2010; Porrello et al., 2011; Senyo et al., 2013). This requirement for pre-existing cardiomyocytes imposes a limit on the extent of damage that the myocardium can sustain and heal through a regenerative response. Using Diphtheria toxin receptor (DTR)-mediated genetic ablation of cardiomyocytes, Wang et al (Wang et al., 2011) reported that at least 40% of the adult zebrafish needs to remain uninjured for regeneration to occur. In neonatal mice, this limit is thought to be even lower, since resection of over 20% of the ventricles results in regenerative failure (Bryant et al., 2015).

In mice, the heart can only regenerate during the first week of life (Porrello et al., 2011). Several phyla and organ systems also display similar age dependent decline of regenerative capacity (Yun, 2015). For example, the tail and limbs of *Xenopus* can regenerate until a refractory period (Beck et al., 2003), and this regenerative ability is abolished post-metamorphosis (Dent, 1962). In insects, imaginal discs and limbs can regenerate during larval and nymphal stages, but not in adults (Nakamura et al., 2008; Smith-Bolton et al., 2009). While we understand that tissues, like the myocardium, lose regenerative ability as they age, we do not know how the competence to regenerate is acquired during development. Understanding how regeneration competence is acquired during development will potentially help us to bestow it to non-regenerative tissues, such as the adult human myocardium.

Transcending the limited ability of vertebrates to regenerate lost parts, experimental animal models have emerged that display remarkable abilities, including whole body regeneration. Champions of regeneration, such Planaria, Cnidarians and Acoels, are found across the phylogenetic tree of metazoans (Tanaka and Reddien, 2011). Unique among them, tunicates are the closest relatives to the vertebrates, offering attractive opportunities to identify regenerative mechanisms built upon a chordate genetic, cellular and molecular toolkit.

How heart regeneration evolved also remains an open question. Early branching vertebrates such as fish and amphibians tend to have higher regenerative capacity (Bely, 2010), pointing to evolutionary losses in mammals, but the evolutionary origins of heart regeneration and state of the vertebrate ancestors remain enigmatic. A true beating heart first evolved in the most common ancestor between the vertebrates and Tunicates (Davidson and Levine, 2003; Kaplan et al., 2015; Stolfi et al., 2010), raising the question as to whether regenerative cardiogenesis evolved in concert or was acquired secondarily.

Tunicate species of the *Ciona* genus are invertebrate chordates, among the closest relatives to vertebrates (Delsuc et al., 2006). Modern evolutionary developmental biological studies using *Ciona* have cast light on the origins of diverse vertebrate traits, including the central and peripheral nervous systems, and the cardiopharyngeal lineage (Abitua et al., 2012; Horie et al., 2018; Stolfi et al., 2010, 2015). By contrast with their inability to compensate for the loss of early fate-restricted lineages in developing embryos (Conklin, 1905), adult and juvenile solitary ascidians readily regenerate distal parts, such as the siphons and neural complex, after complete removal (Dahlberg et al., 2009; Whittaker, 1975; Hirschler, 1914). On the other hand, it is thought that *Ciona* cannot regenerate proximal tissues such as the digestive tract and heart (Hirschler, 1914; Jeffery, 2015a), in part because the animals did not survive the traumatic injuries inflicted upon them. Whether *Ciona* visceral organs could regenerate after more targeted injuries thus remains an open question. Indeed, cardiopharyngeal developmental mechanisms are extensively conserved between *Ciona* and vertebrates (Diogo et al., 2015; Kaplan et al., 2015; Wang et al., 2019), and *Ciona* cardiomyocytes are mononuclear diploid (Millar, 1953), a suggested prerequisite for regeneration (Hirose et al., 2019; Patterson et al., 2017), thus making *Ciona* a prime candidate to revisit its potential for regenerative cardiogenesis. To inflict milder and controlled heart injuries, we adopted the Nitroreductase (NTR) and metronidazole (MTZ)-based inducible chemogenetic ablation system (Curado et al., 2008), and discovered that the *Ciona* heart can regenerate after all. We characterized the cellular and signaling underpinnings of *Ciona* heart regeneration, revealing mechanisms conserved with vertebrates, which point to possibly shared evolutionary origins. Remarkably, we uncovered an unprecedented case of *de novo* regenerative cardiogenesis from endodermal lineages, which challenges established paradigms for heart regeneration. Finally, we discovered that animals acquire their regenerative ability during metamorphosis, illustrating a fundamental transition between deterministic and regulative modes of development.

## Results

### Characterization of the juvenile *Ciona* heart

To study regeneration of the juvenile *Ciona* heart, we first sought live heart markers. The stable transgenic line Tg[MiCiTnIG]2 (Hozumi et al., 2010), hereafter called Mi(TnI>GFP), labels the entire heart and other muscles including the longitudinal muscles (LoM) flanking the heart, atrial siphon muscles (ASM), and oral siphon muscles (OSM) (Figure 1A). We also found that the commonly used monoclonal antibody against myosin heavy chain, MF20 (Bader et al., 1982), specifically labels the cardiomyocytes of the heart, but not the outer layer of the heart, which is referred to as the pericardium (Figure 1A). MF20 does not label siphon and body wall muscles, which supports the idea that these are akin to smooth muscles (Razy-Krajka and Stolfi, 2019).

**Figure 1.**
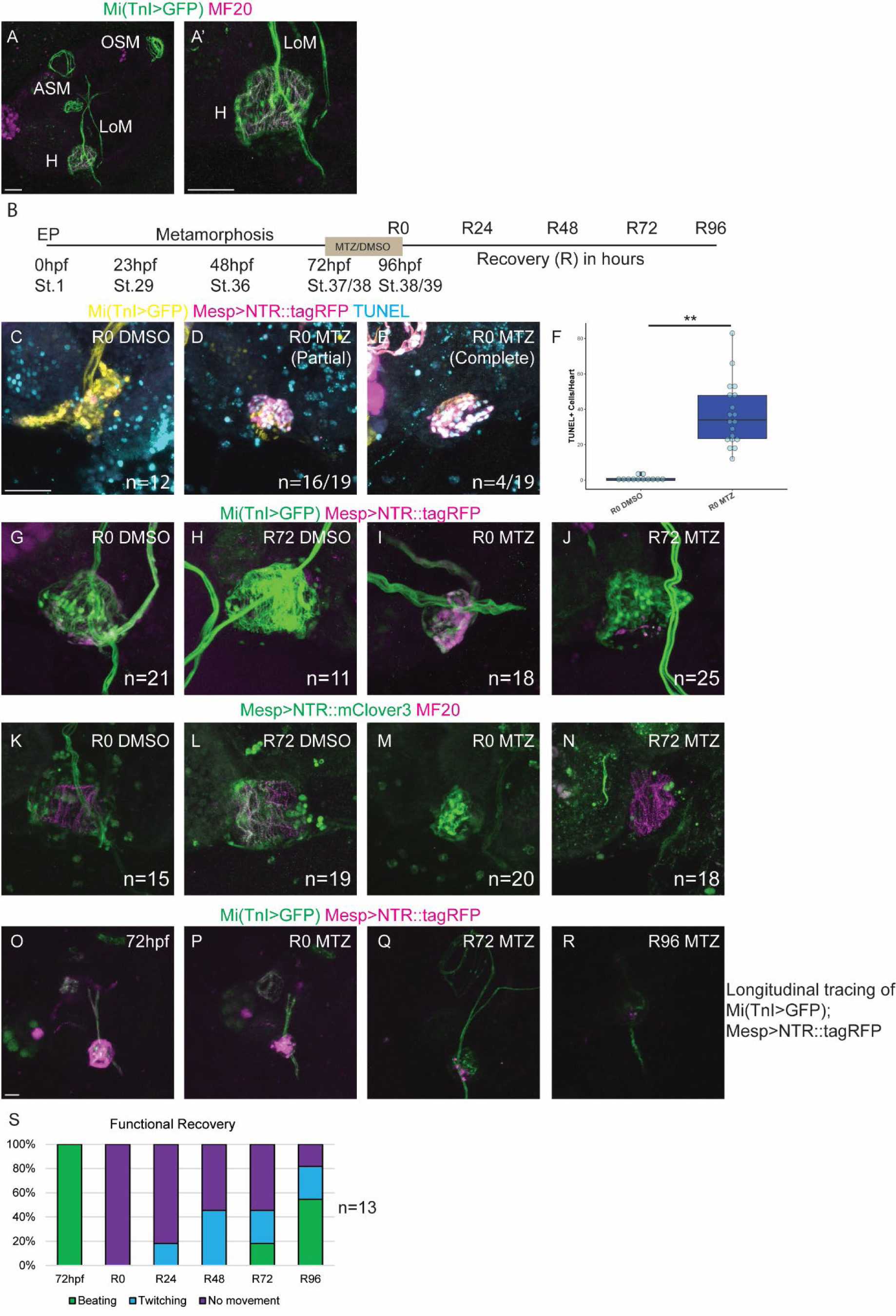
The *Ciona* heart regenerates after extensive ablation of cardiomyocytes. A) 96hpf juvenile *Ciona* transgenic for Mi(TnI>GFP) which labels all muscles including the heart, atrial siphon muscles (ASM), oral siphon muscles (OSM), and longitudinal muscles. MF20 specifically labels the myocardium of the heart (magenta). A’) Visualisation of the heart at a higher magnification highlights that the heart is composed of a TnI>GFP+MF20+ myocardium and a Mi(TnI>GFP)^+^MF20^-^ epicardium. All images with Mi(TnI>GFP) are stained with an anti-GFP antibody to enhance the low levels of expression of the transgene. B) Experimental protocol to induce regeneration in 72hpf (st.37) juveniles. MTZ or 1% DMSO is added to the FASWT for 24 hours. The drug is then washed out. Recovery after treatment in hours is denoted as R#. C-F) TUNEL labeling of Mi(TnI>GFP) juveniles electroporated with Mesp>NTR::tagRFP. C) Undamaged R0 heart treated with DMSO. D) R0 MTZ heart that has a few TUNEL negative cells that survived the ablation protocol, hence referred to as partial ablation. E) R0 MTZ heart that is completely ablated, with every GFP^+^RFP^+^ cell being TUNEL^+^, and no escaping GFP^+^RFP^-^ TUNEL^-^ found. F) Quantification of the number of TUNEL^+^ cells/heart. G-J) Mi(TnI>GFP) hearts electroporated with Mesp>NTR::tagRFP. Looking at undamaged and regenerating hearts at R0 and R72. G) R0 DMSO heart. H) R72 DMSO heart. I) R0 MTZ heart. Note that the heart has collapsed in on itself, with most but not all TnI>GFP^+^ cells being Mesp>NTR::tagRFP^+^. J) R72 MTZ regenerating heart with a greatly recovered number of GFP^+^RFP^-^ cells, with the GFP^+^RFP^+^ cells being progressively cleared and only present as apoptotic debris. K-N) Electroporated with Mesp>NTR::mClover3 and stained with anti-GFP and MF20. K) R0 DMSO shows an undamaged 96hpf heart showing Mesp>NTR::mClover3 being expressed in the entire heart, and MF20 labelling the myocardium. L) R72 DMSO undamaged heart showing largely the same in panel K. M) R0 MTZ. Note that there is a near complete loss of MF20 staining in the dead mClover3^+^ debris. N) R72 MTZ with a restored myocardium as defined by MF20 labeling. Note that there is a few mClover3^+^ debris remaining, but the heart otherwise looks normal. O-R) Longitudinal imaging of the same Mi(TnI>GFP) electroporated with Mesp>NTR::tagRFP juvenile throughout regeneration. O) 72hpf juvenile immediately before MTZ treatment. P) The same juvenile after 24 hours of MTZ treatment, indicated as R0. Note that the heart has collapsed and the apoptotic debris are blebbing. Q) The same juvenile imaged 72 hours later (R72). The apoptotic debris from the Mesp>NTR::tagRFP has been cleared, and GFP^+^RFP^-^ regenerating heart is becoming more and more dominant within the heart. R) The same heart at R96 with even further reduction of NTR::tagRFP^+^ debris and a further increase in Mi(TnI>GFP)^+^ regenerated heart. S) Quantification of functional recovery of MTZ treated hearts every 24 hours. Hearts were scored whether they were beating normally, twitching sporatically (an intermediate state where function is beginning to return), and not beating at all. After damage, the heart stops beating, but eventually regains function over the course of 4 days. Scale bars: 25μm. **p<0.01

### The heart can regenerate in *Ciona* juveniles

The heart of the *Ciona* early juvenile is composed of only 30 to 40 cells, and is difficult to access surgically. To circumvent this issue, we adopted the Nitroreductase (NTR) and Metronidazole (MTZ) genetic ablation system (Curado et al., 2008) to specifically ablate the cardiopharyngeal lineage in a chemically inducible manner. We expressed fluorescent NTR fusion proteins with the *Mesp* enhancer, which is active in B7.5 blastomeres, thus driving NTR in the cardiopharyngeal lineage, including heart cells, and atrial siphon and body wall muscles (Satou et al., 2004; Stolfi et al., 2010). To ablate the heart, we treated *Mesp>NTR::tagFP*-electroporated juveniles with either 10 mM MTZ or DMSO, as carrier control, for 24 hours from stage 37/38 (early juvenile I) at 72 hpf to stage 39 (mid juvenile I) at 96 hpf, before washing out the chemicals and allowing the animals to recover, starting at Recovery time 0h (R0) at 96 hpf (Figure 1B).

We first used a TUNEL assay to label dead cells with fragmented DNA and validate that the NTR/MTZ system can cause inducible and lineage-specific cell death. In control DMSO treated NTR::tagRFP-positive juveniles, we observed sparse to non-existent TUNEL staining in the heart (Figure 1C; Supplemental Figure 1A). By contrast, MTZ-treated animals had numerous TUNEL^+^NTR::tagRFP^+^Mi(TnI>GFP)^+^ cells in the heart, but not in surrounding tissues (Figure 1C-F; Supplemental Figure 1B-C). A subset of hearts were entirely TUNEL positive and visibly collapsed, suggesting that the whole heart was ablated (Figure 1D; Supplemental Figure 1C). We obtained similar results using a Caspase3/7 sensitive fluorescent dye suggesting that these cells are dying from apoptosis (Supplemental Figure 1D-J). Of note, whole heart ablation did not kill the animals, presumably because it is not required for respiratory gas exchange in these early juvenile stages of *Ciona*, and other body wall contractions might suffice to ensure minimal blood movements in the open circulatory system.

This led us to question if the heart can regenerate after this massive amount of cell death. To test this hypothesis, we treated *Mi(TnI>GFP); Mesp>NTR::tagRFP* juveniles with MTZ/DMSO, then fixed and stained them at R0 and after 3-day recovery (R72) (Figure 1G-J). Despite frequent heart collapse following ablation (R0 animals, Figure 1I), they were able to regenerate NTR::tagRFP^-^Mi(TnI>GFP)^+^ cells of the heart, and the amount of NTR::tagRFP^+^ debris appeared to decrease as the amount of GFP^+^RFP^-^ tissue increased (Figure 1J). Similar results were found when staining for the cardiomyocyte specific marker MF20 in *Mesp>NTR::mClover3* damaged hearts (Figure 1K-N). Therefore, the heart is able to regenerate lost tissue after damage. Intriguingly, we are able to see robust regeneration after massive amounts of cell death, with some cases the entire heart appears to have been ablated with little to no surviving cardiomyocytes. This suggests that the entire heart can regenerate *de novo* and provides an opportunity to understand how an animal can regenerate a heart from few to no surviving cardiomyocytes.

To further test and confirm the possibility of heart regeneration, we sought to follow regeneration longitudinally in live animals. We induced heart ablation in Mi(TnI>GFP); Mesp>NTR::tagRFP animals and imaged them in individual glass-bottom wells starting at 72 hpf, and every 24 hours post MTZ washout for four days. We confirmed that NTR-expressing and MTZ-treated hearts first collapsed, progressively forming tagRFP+ debris, while Mi(TnI>GFP)^+^Mesp>NTR::tagRFP^-^ cells emerged and reformed a new heart (Figure 1O-R). Longitudinal tracing of live animals allowed us to examine heart beats as a physiological parameter for functional recovery. Consistent with massive cell death, hearts stopped beating immediately after ablation, but newly formed hearts restored “twitching” first, followed by normal beating by R96 in >50% of the animals (Figure 1S). Taken together, these data indicate that the heart can both regenerate cardiac tissue and restore function in *Ciona* juveniles.

### Cardiomyocyte proliferation contributes to heart regeneration

Next, we sought to characterize the cellular basis for heart regeneration in *Ciona* juveniles. In vertebrates, the heart regenerates via dedifferentiation and proliferation of pre-existing cardiomyocytes, and not from a pool of stem cells (Jopling et al., 2010; Porrello et al., 2011; Senyo et al., 2013). By analogy, we hypothesized that cardiomyocyte proliferation also contributed to heart regeneration in *Ciona*. To assay proliferation, we stained both regenerating and undamaged *Mi(TnI>GFP); Mesp>NTR::tagRFP* hearts for the mitotic marker phospho-Histone H3 (PH3) at 72 hpf (before ablation), R0 (immediately after MTZ/DMSO treatment), and R72 (3 days post MTZ washout). In undamaged DMSO treated hearts, the number of PH3^+^ cardiomyocytes declined over time, while regenerating hearts treated with MTZ from 72hpf-96hpf maintained higher numbers of PH3^+^ cardiomyocytes compared to controls at R0 and R72 (Figure 2A-F). This opens the possibility that increased mitotic activity contributes to replenishing the population of cardiomyocytes during heart regeneration.

**Figure 2.**
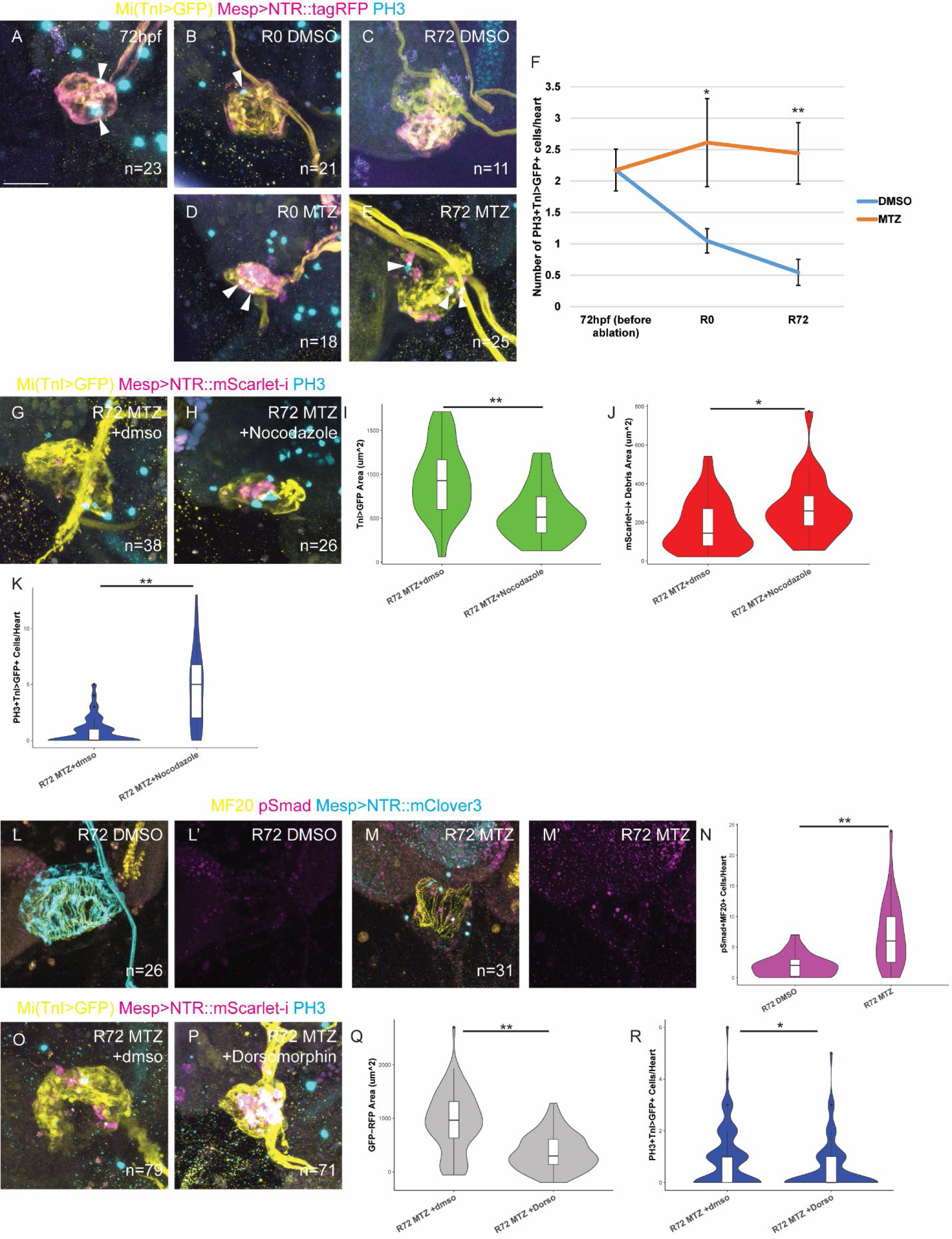
Heart regeneration requires proliferation and BMP signaling. A-E) Mi(TnI>GFP) juveniles at different points during homeostasis or regeneration electroporated with Mesp>NTR::tagRFP stained with anti-GFP to label the heart and anti-PH3 to label mitotic cells. Arrowheads point to PH3^+^TnI>GFP^+^ cells within the heart. A) 72hpf heart with two PH3+ cells. B) R0 DMSO heart with a single PH3^+^ cell. C) R72 DMSO heart without any PH3+ cells within the heart. D) R0 MTZ ablated heart with a couple surviving TnI>GFP^+^ cells also being positive for PH3. E) R72 MTZ regenerated heart with 3 PH3^+^ cells in the heart (arrowheads). F) Quantification of the number of PH3^+^ cells in the heart over time in DMSO treated and MTZ treated animals. The number of PH3^+^ cells decline in undamaged hearts, but in regenerating hearts the numbers increase relative to controls. G-H) To test for the role of cell division during regeneration, 1μg/mL Nocodazole, which arrests cells in mitosis (as evidenced by elevated numbers of PH3^+^ cells) was added to the FASWT (0.1% DMSO as a control) after washing out 10mM MTZ on Mi(TnI>GFP) animals electroporated with Mesp>NTR::mScarlet-I. Incubations with Nocodazole or DMSO was done from R0-R72, which were then fixed and stained for PH3 and GFP. G) Control regenerating heart treated with 0.1% DMSO (R0-R72) after MTZ treatment. H) R72 MTZ heart treated with Nocodazole (R0-R72). Note that it is unable to regenerate and the surviving TnI>GFP^+^ cells are arrested in metaphase, as indicated by PH3. I) Quantification of the average area of TnI>GFP^+^ hearts in each condition. Note that Nocodazole treatment significantly reduces the area of TnI>GFP thus showing that they are unable to regenerate. J) Quantification of the area of Mesp>NTR::mScarlet-I positive debris. Nocodazole treatment increases the amount of debris retained during regeneration. K) Quantification of the number of PH3^+^TnI>GFP^+^ cells per heart. L-M) Mesp>NTR::mClover3 electroporated juveniles stained for pSmad1/5/8, MF20 and GFP in undamaged and regenerating hearts. L) R72 DMSO undamaged heart. M) Regenerating R72 MTZ heart with pSmad^+^ CMs. L’-M’) pSmad staining alone. N) Quantification of the number of pSmad^+^MF20^+^ CMs per heart. O-P) Functional analysis of the role of BMP signaling by treating with either 0.1% DMSO or 10μM Dorsomorphin after MTZ treatment from R0-R72. Mi(TnI>GFP) electroporated with Mesp>NTR::mScarlet-I stained for PH3 and GFP. Q) Quantification of living/regenerating heart area (TnI>GFP area minus NTR::mScarlet-I debris area) in R72 MTZ +0.1% DMSO(R0-R72) versus R72 MTZ +10uM Dorsomorphin(R0-R72). R) Quantification of the number of PH3^+^TnI>GFP^+^ cells per heart in R72 MTZ +0.1% DMSO(R0-R72) versus R72 MTZ +10uM Dorsomorphin(R0-R72). *p<0.05, ** p<0.01, n.s.=not significant. Scale bar: 25μm

Supporting this notion, treatments with low levels of Nocodazole, which blocks cell division by arresting mitotic cells in M-phase, for the entire regenerative period inhibited heart regeneration (Figure 2G, H), as evidenced by a reduction of the Mi(TnI>GFP) area (Figure 2I), and an increase in the amount of Mesp>NTR::mScarlet-i^+^ debris, compared to DMSO-treated controls (Figure 2J). Cells stuck in M phase retained PH3 signals, allowing us to determine that most surviving Mi(TnI>GFP)^+^Mesp>NTR::mScarlet-i^-^ cells were PH3 positive (Figure 2K). This indicated that heart regeneration requires proliferation of an initial population of cardiomyocytes. Of note, because of the mosaic nature of electroporation-based transfection in *Ciona*, our method can yield either partial or complete heart ablation and regeneration. We surmise that, in the case of partial heart regeneration, remaining B7.5 lineage-derived cardiomyocytes provide the initial pool of cells that proliferate and replenish cardiac tissue. On the other hand, complete ablation and whole heart regeneration opens the tantalizing possibility that the initial pool of proliferating cardiomyocytes emerges *de novo* from (a) non-cardiac lineage(s).

### BMP signaling is required for cardiomyocyte proliferation during heart regeneration

A major question that follows the discovery of heart regeneration in *Ciona* is how conserved or divergent the signaling mechanisms are compared to vertebrate heart regeneration. BMP signaling is a conserved signaling pathway found in all Metazoans and is a critical component of the cardiac regenerative response in zebrafish and mice (Wu et al., 2016; Sun et al., 2014; Bongiovanni et al., 2023). To characterize the cellular context of BMP-Smad activity, we first examined patterns of signaling activity by staining for phospho-Smad1/5/8 (pSmad) in undamaged and regenerating hearts labeled with MF20. The number of pSmad^+^MF20^+^ cells was significantly higher in regenerated hearts (Figure 2L-N), suggesting activation during regeneration.

Intriguingly, stomach and intestine cells directly adjacent to the heart also exhibited pSmad staining in both control and regenerative animals. Treatment with the BMP signaling inhibitor Dorsomorphin during regeneration hampered regenerative growth (Figure 2O-Q), and significantly reduced the number of PH3^+^ cardiomyocytes compared to DMSO treated controls (Figure 3R). These data suggest that BMP signaling is a conserved regulator of cardiomyocyte proliferation during heart regeneration in Olfactores.

**Figure 3.**
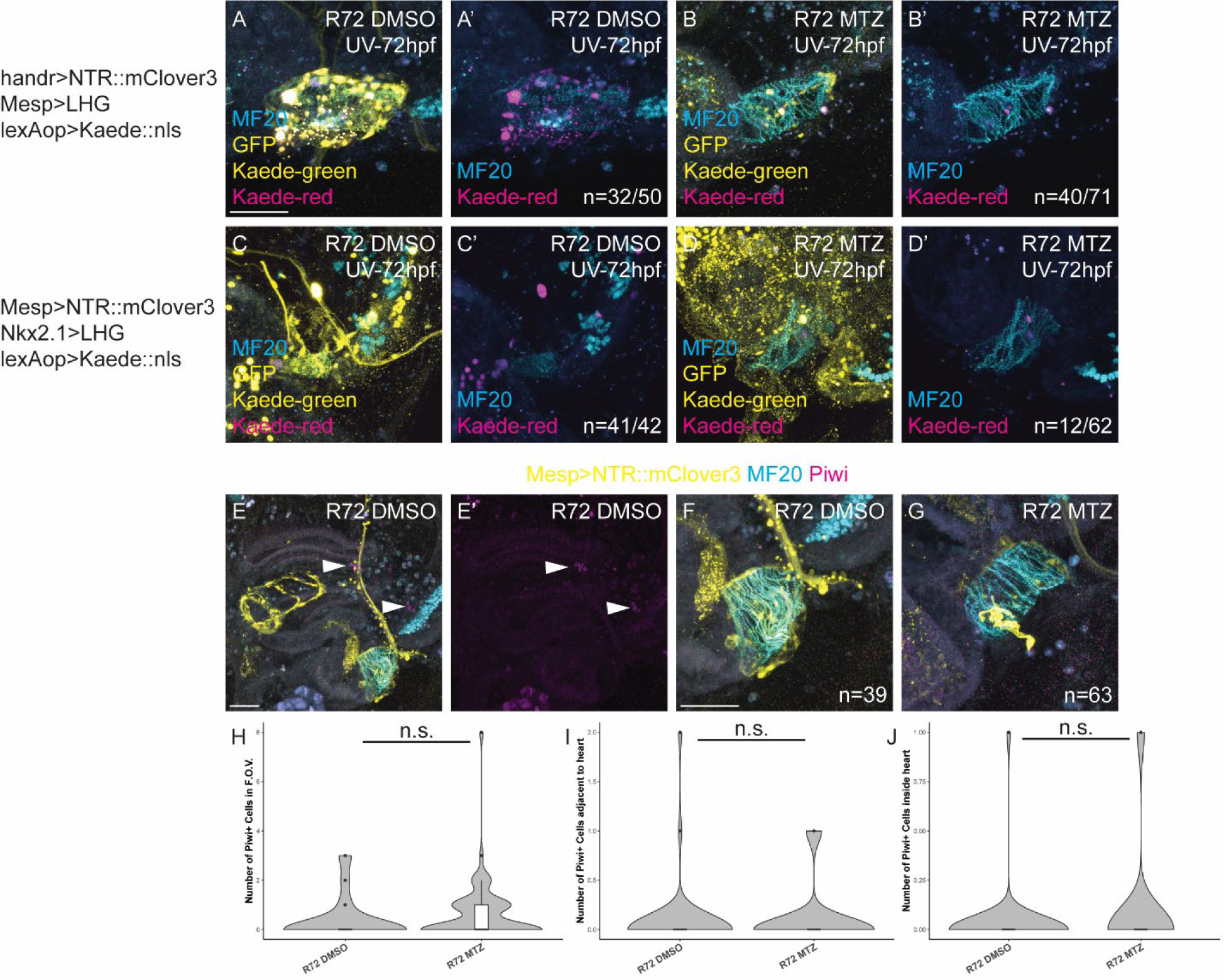
Kaede-based lineage tracing reveals contributions of pre-existing cardiomyocytes and an endodermal source of cardiomyocytes. A-B) Hearts electroportated with Mesp>LHG; lexAop>Kaede::nls and handr>NTR::mClover3. Kaede is converted from green to red fluorescence by treatment with UV illumination on the entire body of the juvenile at 72hpf (beginning of the DMSO/MTZ treatment), then allowed to undergo the normal regeneration protocol until R72. Stained with anti-GFP and MF20. A) R72 DMSO with Kaede-red positive cells within the heart. A’) Kaede-red and MF20 alone without GFP channel. B) Regenerating R72 MTZ heart showing Kaede-red positive cells within the heart. B’) Kaede-red and MF20 alone without GFP channel. C-D) Juveniles electroporated with Mesp>NTR::mClover3, Nkx2.1>LHG, and lexAop>Kaede::nls. Kaede is converted from green to red fluorescence by treatment with UV illumination on the entire body of the juvenile at 72hpf (beginning of the DMSO/MTZ treatment), then allowed to undergo the normal regeneration protocol until R72. Nkx2.1 is expressed in all endoderm-derived cells and is most enriched in the gut. Samples stained for MF20 to label the myocardium and anti-GFP to better visualize Mesp>NTR::mClover3. C) R72 DMSO heart with no Kaede-red positive cells within the heart. C’) Kaede-red and MF20 staining without GFP channel. D) Regenerating R72 MTZ heart showing a Kaede-red positive cell within the MF20^+^ heart. Note that this is a relatively rare event with only 19.7% of all samples showing this. D’) Kaede-red and MF20 staining without GFP channel. E-G) Juveniles electroporated with Mesp>NTR::mClover3 that are either undamaged (E-F) or regenerating (G) stained for Piwi, MF20 and GFP. E) Control juvenile stained for showing a few Piwi^+^ cells near the gill slits and endostyle (arrowheads). E’) Piwi staining alone. F) Same juvenile as (E) at higher magnification. In this field of view (F.O.V) there are no visible Piwi^+^ cells. G) Regenerating heart showing no Piwi^+^ cells near it. H) Quantification of the number of Piwi^+^ cells in the field of view (F.O.V) near the heart. I) Quantification of the number of Piwi^+^ cells directly adjacent to the heart. J) Quantification of the number of Piwi^+^ cells inside the MF20^+^ heart. n.s.=not significant. Scale bar 25μm.

### A Small-Scale Inhibitor Screen for Additional Conserved Signaling Pathways Required for Heart Regeneration

Encouraged by our BMP signaling results, we surveyed other druggable signaling pathways that may exert conserved roles in heart regeneration. We focused on JAK/STAT, Hippo/Yap, and Hedgehog signaling pathways, which are all essential for cardiac regeneration in vertebrates (Choi et al., 2013; Fang et al., 2013; Kawagishi et al., 2018; Leach et al., 2017; Monroe et al., 2019; Wang et al., 2015; Xin et al., 2013). We treated animals with inhibitors immediately after MTZ washout and allowed them to recover for 3 days before staining for PH3 and either GFP for *Mi(TnI>GFP)* transgenic animals or MF20 staining, and assessed for the number of PH3 positive cardiomyocytes and the size of the regenerated area. We found that inhibitors targeting the JAK/STAT and Hippo/Yap pathways resulted in impaired regeneration, but we did not find any phenotype upon inhibition of hedgehog signaling (Supplemental Figure S2). This led us to propose that BMP/Smad, JAK/STAT and Hippo/Yap pathways form a conserved signaling cocktail required for heart regeneration in olfactores.

### Kaede-based lineage tracing reveals contributions of pre-existing cardiomyocytes and an endodermal source of cardiomyocytes

The *Ciona* heart can regenerate after extensive damage. We noticed that a subset of animals display damaged hearts entirely labeled with TUNEL, suggesting complete ablation (Figure 1D). Such whole heart regeneration implies the existence of (an) alternative source(s) of cells that contribute(s) to regenerating cardiomyocytes. We sought to prospectively trace embryonic lineages that contribute to regenerating cardiomyocytes, using UV-induced green-to-red photoconversion of the fluorescent protein Kaede (Ando et al., 2002), which is commonly used in *Ciona* for lineage tracing through metamorphosis (Horie et al., 2011; Nakazawa et al., 2013; Razy-Krajka et al., 2014; Wang et al., 2019). Since vertebrate hearts regenerate via dedifferentiation and proliferation of pre-existing cardiomyocytes (Jopling et al., 2010; Kikuchi et al., 2010; Porrello et al., 2011; Senyo et al., 2013), we first tested if the heart regenerates from pre-existing cardiomyocytes following partial ablation. We used the *Hand-r*/*NoTrlc* cardiopharyngeal enhancer (Woznica et al., 2012), which is more mosaically inherited and thus favors partial ablation, to express NTR::mClover3 alongside *Mesp>LHG* and *lexAop>Kaede::nls* since the the lexAop-LHG system enhances expression compared to Mesp>Kaede::nls alone (Tolkin and Christiaen, 2016). We used UV light to photoconvert Kaede from green to red at 72 hpf while starting MTZ or DMSO treatments. We allowed animals to recover for 72 hours following 24 hrs treatments, fixed and stained for MF20 to label the myocardium. We found Mesp>>Kaede-Red^+^ cells in both the control and regenerating hearts (Figure 4A-B). This observation is consistent with persistence and a contribution of pre-existing cardiomyocytes to the regenerated heart following partial damage.

**Figure 4.**
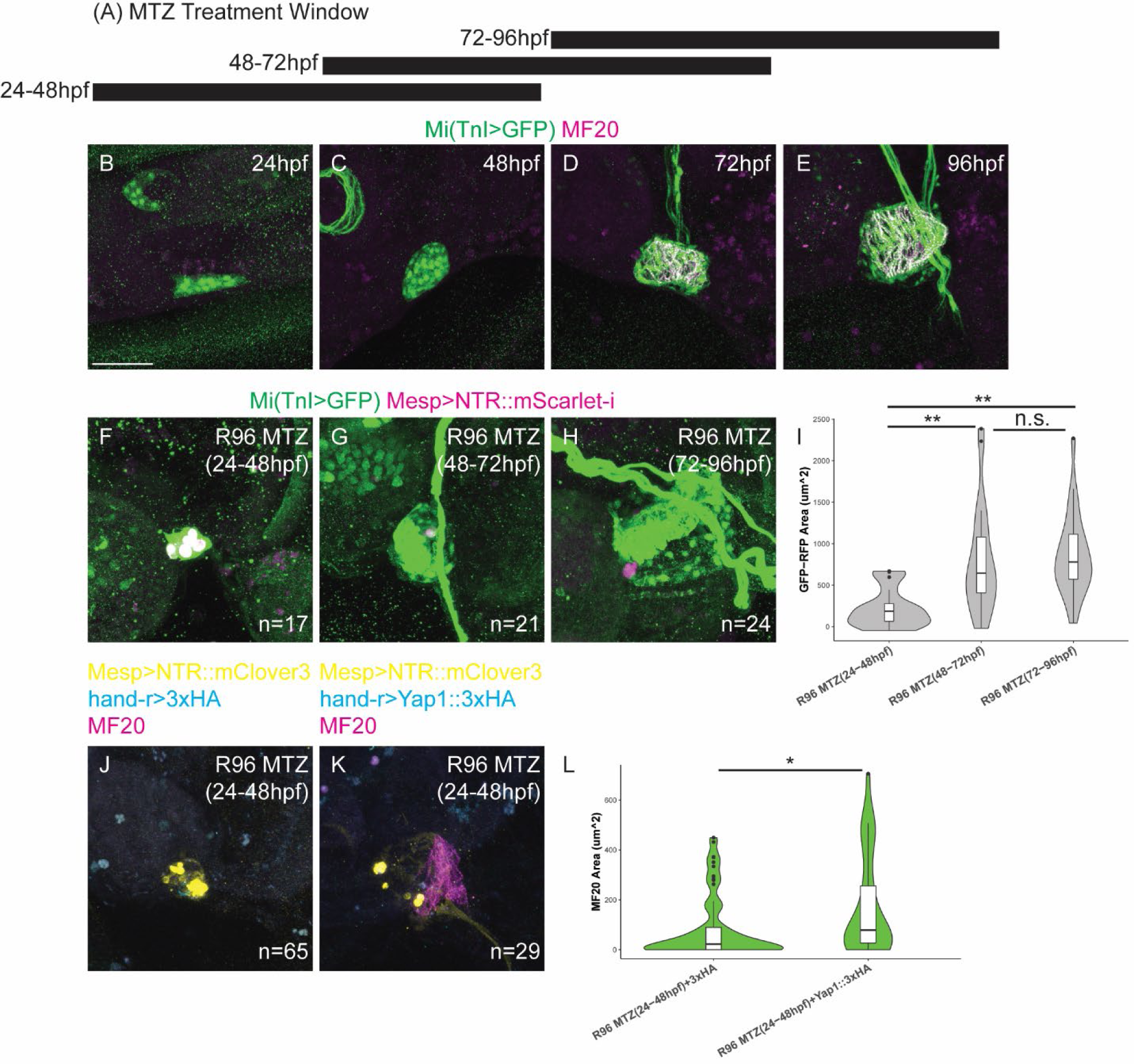
Regenerative capacity is acquired post-embryonically. A) Experimental design showing the treatment window of MTZ treatments throughout development, with the corresponding stage under the bars. B-E) Developmental time course of the heart through metamorphic development sampled every 24 hours. Mi(TnI>GFP) animals were stained for anti-GFP and MF20 to highlight the developmental maturation of the heart throughout metamorphosis. B) 24hpf larva. No MF20 staining found in the heart (wedge shaped structure). C) 48 hpf mid to late body axis rotation stage juvenile. The heart has formed a ball shape but is still undifferentiated as it still does not have MF20 labeling. D) 72hpf juvenile with clear cardiomyocyte differentiation as shown with MF20 labeling. E) 96 hpf juvenile. The heart has even more clearly defined myocardium and a separate epicardium is also evident. F) Mi(TnI>GFP) juvenile electroporated with Mesp>NTR::mScarlet-I that was treated with MTZ from 24-48 hpf (st.29 late swimming larvae to st.36 late body axis rotation) then allowed to recover until R96. These hearts failed to regenerate. G) Mi(TnI>GFP) juvenile electroporated with Mesp>NTR::mScarlet-I that was treated with MTZ from 48-72 hpf (st.36 late body axis rotation to st.37/38 early juvenile I/II) then allowed to recover until R96. Hearts regenerated robustly. H) Mi(TnI>GFP) juvenile electroporated with Mesp>NTR::mScarlet-I that was treated with MTZ from 72-96hpf (st.38 early juvenile II) then allowed to recover until R96. Hearts regenerated robustly. I) Quantification of living/regenerating heart area (TnI>GFP area minus NTR::mScarlet-I debris area). R96 MTZ(24-48 hpf) are significantly reduced compared to the later ablated hearts that can regenerate. J) Mesp>NTR::mClover3; hand-r>3xHA electroporated juvenile treated with MTZ from 24-48 hpf and then allowed to recover until R96. Stained for MF20, GFP and HA. These animals fail to regenerate. K) Mesp>NTR::mClover3; hand-r>Yap1::3xHA electroporated juvenile treated with MTZ from 24-48 hpf and then allowed to recover until R96. Stained for MF20, GFP and HA. These animals are able to partially regenerate the myocardium stained for MF20. L) Quantification of MF20 staining area. * p<0.05 ** p<0.01 n.s.=not significant. Scale bar 25μm.

We hypothesized that, following complete ablation, cardiomyocytes likely emerge from an extracardiac source of cells. The juvenile heart is surrounded by endoderm-derived tissues, including the intestine, stomach, endostyle and retropharyngeal band, which we saw as likely candidate source(s) of cells for *de novo* cardiogenesis during regeneration. We used the early endoderm marker *Nkx2.1>LHG; lexAop>Kaede::nls* (Ristoratore, 1999), to trace cells following induced damage with *Mesp>NTR::mClover3*, which increases the likelihood of complete ablation. We converted Kaede at 72 hpf and treated the animals with MTZ/DMSO until 96 hpf, then allowed them to recover for 3 days. We fixed and stained for MF20 to label the myocardium. Strikingly, we found Nkx2.1>>Kaede::nls^+^ cells in ∼19% of the regenerated, but not control hearts (Figure 4C-D). We observe few Kaede^+^ cells in these regenerated hearts, possibly due to mosaic inheritance of transgenes, and/or dilution of the Kaede proteins through proliferation over time. This suggests that cells from an endodermal origin can contribute cardiomyocytes to regenerating hearts following complete ablation.

A population of Piwi^+^ progenitors has been proposed to function as a regenerative cell source of post amputation in *Ciona* (Jeffery, 2015b, 2018). Based on their alkaline phosphatase activity (Jeffery 2015b), these cells are likely endodermal in origin. We considered the possibility that these same Piwi^+^ progenitors could serve as this alternative cell source to regenerate the heart. We stained for Piwi using a previously characterized antibody shown to work in *Ciona* (Jeffery, 2015b) in damaged and regenerating juveniles. We found few Piwi^+^ cells in both conditions, and they were rarely found in and around the heart (Figure 3E-J). Their distribution appeared random and did not appear to target the damaged heart. This leads us to conclude that Piwi^+^ progenitors are not likely to be the endodermal cell source that can regenerate the heart after complete ablation.

### Regenerative capacity is acquired post-embryonically

Ascidians have long been models of mosaic development (Conklin, 1905), where each blastomere produces a defined lineage in an irreplaceable manner, by contrast with regulative early development in vertebrates. For instance, blocking specification of cardiac progenitors in early *Ciona* embryos inhibited heart formation in juveniles, illustrating the animal’s inability to compensate for the absence of cardiogenic lineage during embryogenesis (Satou et al., 2004). Ablation of blastomeres in the embryo also results in a complete loss of that corresponding lineage. We thus sought to determine when *Ciona* acquires the ability to regenerate the heart, a fundamental transition between deterministic and regulative development. We performed ablations at different developmental stages, then allowed them to recover for 4 days and assayed heart regeneration. MTZ takes 20 to 24 hours to induce signs of apoptosis (Supplemental Figure 1D-K), limiting temporal resolution to 24-hour increments (Figure 4A-E). Early juveniles ablated from 48-72 hpf achieved heart regeneration to a degree comparable to positive control juveniles that were ablated from 72-96 hpf (Figure 4G, H). By contrast, 24 hpf settled larvae treated with MTZ until 48 hpf (late body axis rotation) failed to clear debris and regenerate their hearts (Figure 4F,I). Accordingly, the amount of regenerated heart tissue, assayed as the area of Mi(TnI>GFP) positive, Mesp>NTR::mScarlet-i negative tissue, was significantly lower in animals treated from 24-48 hpf (Figure 4I). Therefore, *Ciona* acquires heart regeneration capacity between 48hpf and 72hpf, which correspond to late stages of metamorphosis (stages 36 to 37, (Hotta et al., 2020)) and the expression of cardiac myosin (Figure 4B-E). How do non-regenerative larvae acquire regenerative competence through metamorphosis? We sought to test candidate pro-regenerative factors that can bestow regenerative competence to non-regenerative stages of *Ciona* when overexpressed in the cardiopharyngeal lineage. The Hippo signaling co-transcription factor Yes Associated Protein (Yap) promotes reprogramming of adult mammalian cardiomyocytes to a more regenerative state (Monroe et al., 2019). We therefore considered Yap1 as a candidate regulator for the regenerative transition. For instance, Yap1 over-expression in swimming larvae sufficed to increase the MF20-positive area following ablation, suggesting that Yap1 can bestow regenerative competence upon previously non-regenerative cells (Figure 4I-K).

## Discussion

In *Ciona*, amputation studies where only proximal fragments containing visceral organs can regenerate distal structures such as the brain and siphons have led to the traditional assumption that the *Ciona* heart cannot regenerate (Hirschler, 1914; Jeffery, 2015a; Whittaker, 1975). Challenging these assumptions, here we used NTR/MTZ-based ablation of the cardiopharyngeal lineage, and showed that the *Ciona* heart can actually regenerate after damage and,in some cases, through *de novo* cardiogenesis from (an) extra cardiac source(s), an unprecedented finding when compared to the vertebrates.

Colonial tunicates such as *Botryllus schlosseri* can undergo whole body regeneration (Kassmer et al., 2016), thus regenerating heart tissue among other organs. However, colonial tunicates undergo non-embryonic cardiogenesis as part of their asexual reproduction cycles through budding/blastogenesis. *De novo* cardiogenesis in colonial ascidians could be viewed as homeostasis, rather than a regenerative response. By contrast, since *Ciona* only reproduces sexually, regenerative programs are deployed in response to injury.

Vertebrate hearts regenerate via dedifferentiation and proliferation of pre-existing cardiomyocytes (Jopling et al., 2 010; Kikuchi et al., 2010; Porrello et al., 2011; Senyo et al., 2013) and there are no cardiogenic adult stem cells. By contrast, the *Ciona* heart could regenerate the entire heart after complete ablation, and appears to have an endodermal cell population that contributes to the new myocardium. It was recently found that promoting the development of both cardiac and intestinal organoids simultaneously results in an enhanced maturation of the cardiomyocytes (Silva et al., 2021), suggesting that the endoderm-derived cells are required for proper cardiac development. It has been suggested that the oral siphon and neural complex regenerates via a population of Piwi^+^ stem cells found in the branchial basket (Jeffery, 2015b, 2018). We believe these are not derived from Piwi^+^ stem cells because we do not observe Piwi^+^ cells to be within or around the heart during regeneration more than we see in controls, and that the distribution of these cells appear random and may just be a subset of hemocytes that are passively circulating (Figure 3E-J). Indeed, Piwi^+^ cells are very rare in the stages we’ve examined, which suggests that they may not provide any regenerative function.

There is other evidence for alternative cell sources during regeneration when the primary regenerative cell source is depleted, another cell population contributes to the regenerating structure to ensure that function is restored. For example, when pancreatic β cells are depleted, *α* cells can transdifferentiate into β cells and restore the damaged pancreas’s ability to produce insulin (Thorel et al., 2010). In the regenerating zebrafish caudal fin, when all the osteoblasts are ablated an alternative cell source contributes to bone regeneration instead of the pre-existing osteoblast population (Singh et al., 2012). In the regenerating intestine, if the Lgr5^+^ stem cells are ablating, the transit amplifying cells can dedifferentiate to a Lgr5^+^ stem cell and become competent to self-renew and allow the intestine to undergo homeostatic regeneration (Tetteh et al., 2016). There is also a wealth of *in vitro* models where fibroblast-derived induced pluripotent stem cells can be coaxed to differentiate to cardiomyocyte-like cells (Takahashi et al., 2007). Out of all these examples, the transdifferentiation is mediated by cells derived from the same germ layer. What makes the gut to heart conversion unique is that the heart and gut are derived from different germ layers. It is possible that this endoderm to cardiac fate conversion could also occur in mammals or zebrafish, as researchers do not look at two organs at the same time—researchers usually remove the organ of interest before further processing so we ignore inter-organ dynamics. Another possibility why this might not be possible in vertebrates is that while *Ciona* juveniles can survive around a week without a heart (Satou et al., 2004), severe damage to a vertebrate heart result in a non-regenerative scar and/or lethality (Wang et al., 2011). It is also possible that this is a tunicate-specific adaptation and is absent in vertebrates. It will be essential to determine the mechanisms of how an endodermal cell can transdifferentiate to a cardiomyocyte *in vivo* using the power of *Ciona*.

The highly determinative or “mosaic” development of tunicates has been appreciated for over a hundred years (Conklin, 1905), where ablation of individual lineages result in no compensatory regeneration of those lineages. Despite this rigid form of development up to the larval stage, many adult ascidians can regenerate after amputation (Dahlberg et al., 2009; Kassmer et al., 2016; Shenkar and Gordon, 2015; Whittaker, 1975). Despite this being common knowledge in the field, no one has directly tested what are the mechanisms of this developmental acquisition of regenerative capacity. We found that the mosaic to regenerative transition occurs sometime between day 2 and day 3 of development, which is roughly in the late body rotation and early juvenile I stages (Hotta et al., 2020). Intriguingly, this correlates with the maturation of the heart and the onset of heartbeat. This suggests that a tissue needs to be mature enough to be able to regenerate, as opposed to the current paradigm where it is suggested that immature tissues regenerate better than fully mature tissue. We also discovered that Yap1 is able to reprogram regeneration-incompetent heart cells in the larval stage and early metamorphic stages to a regeneration-competent tissue, much like what was discovered in mice (Monroe et al., 2019). We also note that NTR is rather slow at ablating tissues, so improved methods of ablating cardiomyocytes will be necessary to refine the transition to a more distinct point in time in metamorphic development. Future work will be focused on further refining when the transition occurs using faster methods of ablation and identifying novel factors and/or cell types that are necessary and sufficient for the acquisition of regenerative competence.

Heart regeneration was previously thought to be solely in the realm of vertebrate regenerative abilities. The discovery that *Ciona* can robustly regenerate the heart points to an ancient origin of cardiac regeneration prior to the split between vertebrates and tunicates. We also found conserved signaling pathways, such as BMP/Smad signaling, that are required for regeneration. This suggests that many of the known signaling pathways are conserved in controlling the cardiac regenerative response across Olfactores. Given that cephalochordates do not have a heart and just a contractile vessel (Moller and Philpott, 1973; Pascual-Anaya et al., 2013; Simõ es-Costa et al., 2004), this suggests that the heart had a robust regenerative capacity at the origin of the beating heart itself, or was acquired shortly after. The heart is thought to have evolved from blood vessels that became contractile (Simõ es-Costa et al., 2004), and vertebrate blood vessels are highly regenerative (Evans et al., 2021); we hypothesize that the heart may have inherited its regenerative capacity from the blood vessel regenerative program, but this remains to be directly tested. Finally, the exciting recent discovery of a fossil tunicate that had the same basic body plan as *Ciona* and has evidence of cardiopharyngeal lineage derived longitudinal muscles (Stolfi et al., 2010) from 500 million years ago (Nanglu et al., 2023) suggests how deeply ancient heart regeneration may be.

In conclusion, we have discovered that *Ciona* can indeed regenerate its heart after damage, and it is an exciting emerging model that will be used to identify the cellular, developmental, and evolutionary origins of cardiac regeneration.

## Materials and Methods

### Animals

Wild type *Ciona robusta* (formerly known as *Ciona intestinalis* type A) was obtained from M-REP. The stable transgenic line *Mi(TnI>GFP)* was maintained in our live rearing facility for multiple generations as previously described (Ohta et al., 2020). Briefly, sperm from the Mi(TnI>GFP) was used to fertilize wild WT eggs. The resulting larvae were allowed to settle on uncoated petri dishes and underwent metamorphosis. At 5 days post fertilization, the juveniles were fed a 1:10 dilution of *Isochrysis galbana* and *Chaetoceros gracilis* in Filtered Artificial Seawater buffered with TAPS (FASWT) every other day for 2-3 months before harvesting sperm from the resulting adults to use for experiments or for maintaining further generations.

### Constructs and Electroporation

NTR was PCR amplified from the *pCS2-Nfs.B* construct (Horstick et al., 2015). NTR was then digested with NotI and SpeI and inserted into the *Mesp>tagRFP* construct, creating *Mesp>NTR::tagRFP*. In order to tag NTR with other tags, standard subcloning with restriction enzymes was performed to create *Mesp>NTR::mScarlet-i* and *Mesp>NTR::mClover3*. To generate *lexAop>Kaede::nls*, the *Mesp>Kaede::nls* (Razy-Krajka et al., 2014) was digested with AscI and NotI to remove the *Mesp>* enhancer. The enhancer was replaced with the *hand-r>* and *lexAop>* enhancers. *Nkx2.1>LHG* was previously described (Tolkin and Christiaen, 2016). The *Yap1* ORF KY21.Chr7.860.v1.SL1-1 was amplified from cDNA isolated from 96hpf juveniles and cloned via TOPO-TA cloning. *Yap1* was then amplified from the *TOPO-Yap1* vector and inserted into *hand-r>3xHA* using NotI and SpeI.

Electroporations were performed as previously described (Christiaen et al., 2009). For the NTR containing constructs, 70ug of DNA was used. For Nkx2.1>LHG and Mesp>LHG, 30μg was used. *lexAop>Kaede::nls* was used at 50μg. *hand-r>3xHA* and *hand-r>Yap1::3xHA* were used at 70μg.

### Heart ablation assay

Wild type or Mi(TnI>GFP)/+ zygotes were electroporated with 70ug of the Mesp>NTR::tag construct, in addition to any other construct mentioned above at the corresponding concentration. The resulting electroporated embryos were allowed to grow up to the larval stage at 23-24hpf at 18°C. Larvae were transferred to an uncoated petri dish with FASWT and allowed to settle. 10mM KCl was used to induce a synchronized metamorphosis before washing out with FASWT one day later. At 72hpf, juveniles were treated with 10mM Metronidazole (Sigma) dissolved in DMSO, or 1% DMSO as an undamaged control in FASWT for 24 hours in the dark. After the ablation period, the MTZ/DMSO+FASWT solution was washed out for fresh FASWT and allowed to recover. Importantly, due to the mosaic nature of the electroporation method, ablation was confirmed in regenerating hearts by checking for NTR::FP fluorescence in the cellular debris within and around the heart (defined by Mi(TnI>GFP) fluorescence or MF20 staining). Animals without any evidence of cellular debris at any stage were excluded from any analysis as there was no evidence for ablation. DMSO treated hearts were only imaged if the electroporated transgene was visible; usually undamaged fluorescence was dimmer than in the debris. For the regenerative transition mapping experiments, MTZ/DMSO treatments were done from 24-48hpf, 48-72hpf and 72-96hpf. All experiments were performed at 18°C.

### Immunofluorescence

Animals at the proper stage were anesthetized in 300μg/mL MS-222 in FASWT for 5 minutes before removing them from their attachments on the petri dish. Animals were then fixed in MEM-PFA (4% PFA, 100mM MOPS pH 7.4, 500mM NaCl, 1mM EGTA, 2mM MgSO_4_, 0.05% Tween-20) at room temperature for 30 min. Following two washes with 1xPBS samples were either processed directly or were dehydrated in a graded ethanol series and stored in −20°C overnight or longer before rehydrating in a PBT/Ethanol series. MF20 labeling requires this dehydration-rehydration step. Samples were washed thrice in 1xPBS containing 0.1% Tween-20 (PBT) before digesting with 5-10μg Proteinase K for 25 min at 37°C. ProK treatment was then quenched in 2mM Glycine in PBT and two further PBT washes before being post-fixed in 4%PFA in PBT for 20 min. After a wash in PBT, juveniles were cleared in three 20-minute washes with 50mM NH_4_Cl in PBT. Juveniles were further permeabilized in 1xPBS containing 0.1% Tween-20 and 0.25% TritonX-100 (PBTTx) three times 10 minutes per wash. Samples were then blocked with 3% BSA in PBT for 30 minutes before incubating with primary antibody containing solution. Primary antibodies were diluted in 2% Normal Goat Serum (NGS) in PBT and were incubated with the samples overnight at 4oC. After primary antibody incubation, samples were washed three times with PBT then incubated overnight at 4°C with secondary antibodies diluted in 2% NGS in PBT. After secondary antibody incubation, samples were washed three times in PBT and were transferred into Prolong Gold Antifade reagent (Thermo-Fisher Scientific) which required an overnight equilibration step before mounting on slides flanked with double sided tape as spacers and sealed with clear nail polish.

Antibodies used were rabbit anti-GFP ab290 (Abcam) 1:1000, chicken anti-GFP (Aves Labs) 1:1000, mouse anti-MF20 (Developmental Studies Hybridoma Bank) 1:100, rabbit anti-pSmad1/5/8 41D10 (Cell Signaling Technology) 1:250, rabbit anti-PH3 (Cell Signaling Technology) 1:500, mouse anti-PH3 (Cell Signaling Technology) 1:500, rabbit anti-Piwi ab5207 (Abcam) 1:200, and rat anti-HA (Roche) 1:250. AlexaFluor secondary antibodies were used at 1:500 dilution. Samples were imaged on a Leica SP8 X Confocal Microscope.

The Click-iT TUNEL assay (Thermo-Fisher Scientific) was performed following the manufacturer guidelines but replaced all steps calling for PBS with PBT. CellEvent Caspase3/7 Green detection reagent (Thermo-Fisher Scientific) was used to label cells with caspase activity 30 minutes prior to fixation. A 1:500 dilution was used with living juveniles suspended in a 1.5 mL microcentrifuge tube. After fixation, multiple washes in PBT were performed and NH_4_Cl in PBT was used to clear before mounting in Prolong Gold Antifade supplemented with DAPI.

### Longitudinal tracing of regenerating hearts

72 hpf Mi(TnI>GFP) animals with beating hearts were selected for fluorescence for expressing the Mesp>NTR::tagRFP construct and placed individually inside a glass bottom Lab-Tek II Chambered #1.5 German coverglass system (Fisher Scientific) containing MTZ in FASWT. Animals were imaged on a Leica SP8 confocal every 24 hours pre and post MTZ treatment. The MTZ in FASWT was washed out carefully to keep the juveniles within the wells at R0 and replaced with fresh FASWT that would remain for the rest of the experiment. Scoring for heartbeat was performed during the same imaging session. Animals were scored for either having no heartbeat, twitching (irregular movement of heart muscle), or normal heartbeat (smooth regular beating motion).

### Kaede photoconversion

Whole 72hpf animals electroporated with Kaede::nls containing constructs for lineage tracing analysis were converted using UV (405 nM) light from either a fluorescent microscope or the OptoWELL system (Opto Biolabs). Kaede conversion took 5-10 seconds per juvenile on the microscope or 1 minute on the OptoWELL for all juveniles at once. Juveniles were then transferred back into the plates containing FASWT+MTZ/DMSO and progressed until the desired stage for further analysis.

### Inhibitor treatments

Inhibitors were all dissolved in DMSO to make a stock solution. The stock solution was then diluted 1:1000 in FASWT then added to the plate containing juveniles. An equivalent dilution of DMSO was used as a control. Drugs were always added after the ablation period immediately following MTZ washout, unless otherwise stated. The drug+FASWT containing solution was exchanged for fresh drug every other day until fixation. The working concentration for each drug was as follows: Nocodazole (Sigma Aldrich) 1ng/mL Dorsomorphin (Selleckchem) 10μM, Ruxolitinib (Selleckchem) 10μM, Vismodigeb (Selleckchem) 10μM, and Verteporfin (Tocris Biosciences) 250nM. All inhibitor experiments were performed in the dark.

### Image Quantifications

All images were processed using FIJI Software (ImageJ; NIH). The *Ciona* juvenile heart is a single layered epithelium and therefore can be approximated in two dimensions on a maximum intensity projection. To quantify the size of the heart the freehand selection tool was used to trace around the heart as defined by Mi(TnI>GFP) or MF20 and took the area calculation using the measure function. When necessary, this was also used to take the fluorescence intensity of the ROI. To count the number of pSmad1/5/8^+^ or PH3^+^ cells in the heart, manual counting using the multipoint tool on the z-stack was used. pSmad^+^ or PH3^+^ cells were only counted if they colocalized with GFP or MF20.

### Statistics

Statistical analysis was performed by Rstudio and Excel. Error bars are represented as standard error of the mean. Significance was determined by Welch’s t-test. The investigators were not blinded, nor was sample size predetermined.

## Author Contributions

K.J.S conceived the study. L.C. obtained funding. K.J.S. and L.C. designed experiments. K.J.S. performed experiments, conducted analyses, and prepared figures. K.J.S. and L.C. wrote the paper.

## Acknowledgements

The authors would like to thank the Christiaen lab for the helpful comments and suggestions on the project. We would like to thank Yasunori Sasakura for providing the Mi(TnI>GFP) transgenic sperm that established the transgenic *Ciona* colony. This work was supported by Trans-Atlantic Network of Excellence award from the Leducq Foundation 15CVD01 to L.C., and by NIH awards R01 GM096032, R01 HD096770 and R01 HL108643 to L.C., and by core funding from the Michael Sars Centre, UiB to L.C.

