## Supplemental figures and legends for "The Chordate Origins of Heart Regeneration"

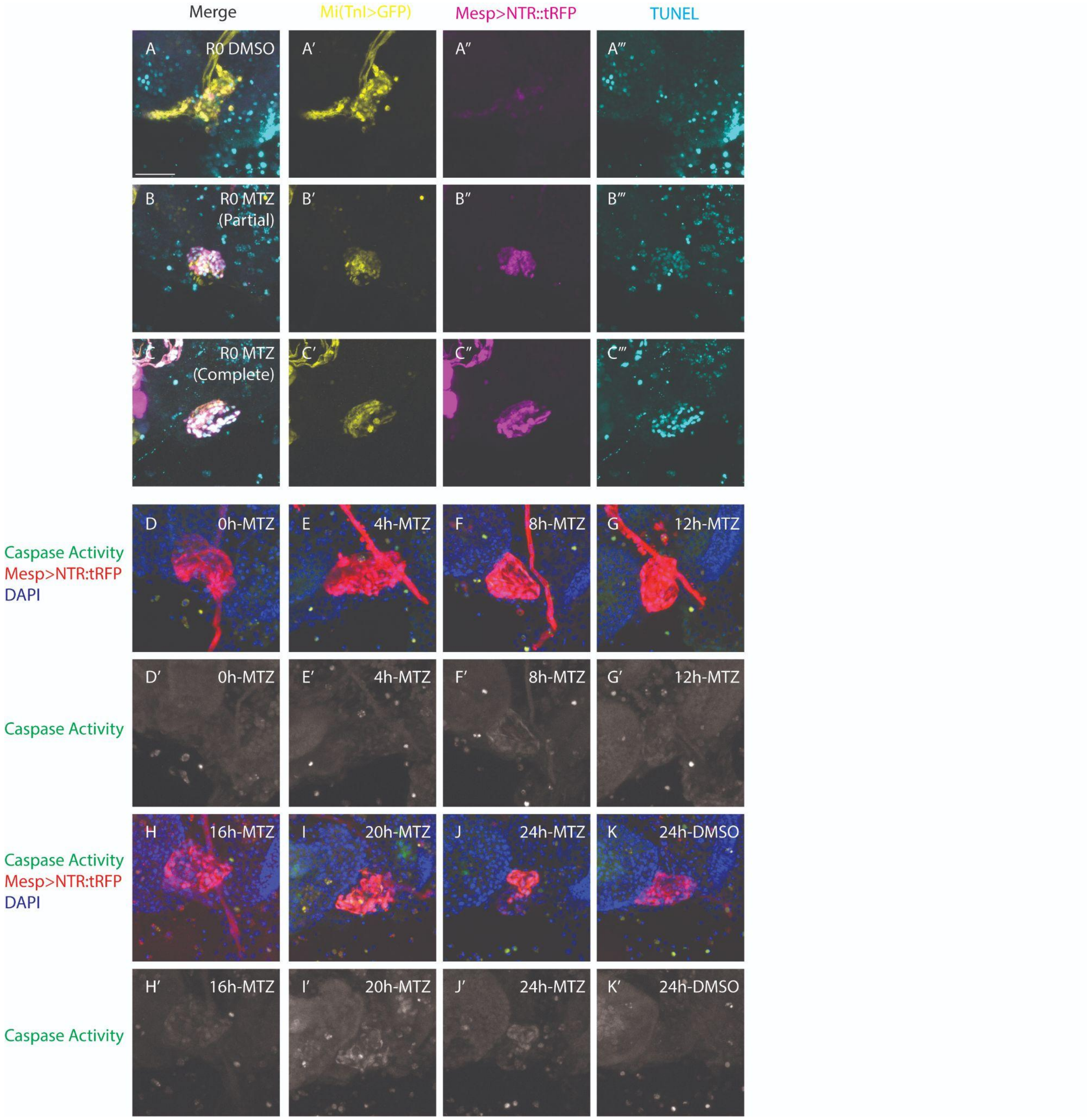


**Supplemental Figure 1. Characterization of Cell Death in ablating *Ciona* hearts.** A-C) TUNEL labelling experiments in DMSO and MTZ treated hearts from Figure 1 in the main text with separate images per channel. A'-C') Mi(TnI>GFP). A''-C'') Mesp>NTR::tagRFP. A'''-C''') TUNEL. D-J) Mesp>NTR::tagRFP treated with MTZ and fixed every 4 hours (starting at 0h and ending at 24h) and stained for CasEvent3/7 prior to fixation to visualize caspase activity in apoptotic cells. It takes about 20-24 hours for reliable detection of both caspase activity and the collapsed heart phenotype of apoptotic debris. K) Mesp>NTR::tagRFP DMSO control heart with no caspase activity in the heart. D'-K') Caspase Activity alone. Scale bar: 25µm

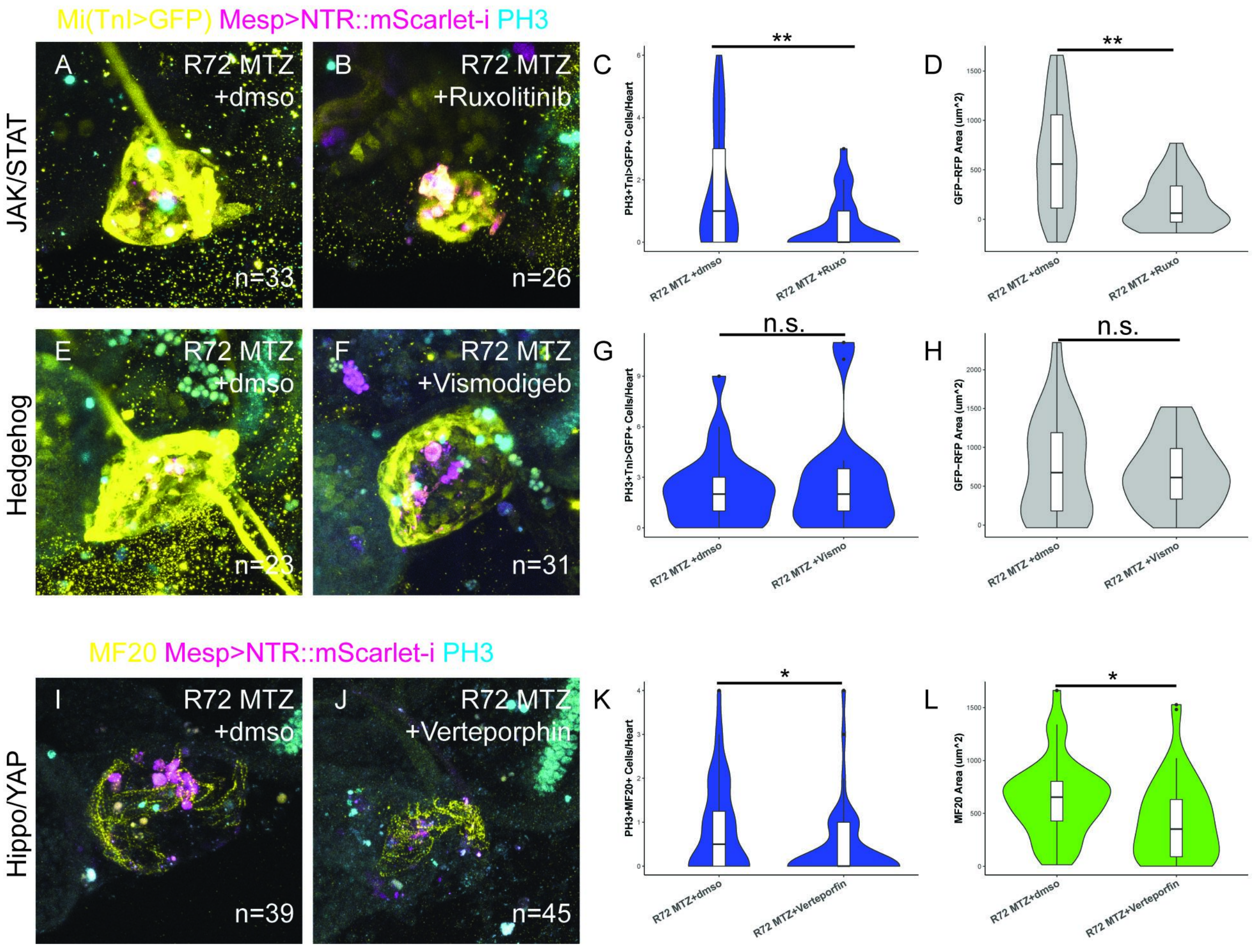


**Supplemental Figure 2. Pharmacological mini-screen for signaling pathways required for heart regeneration.** A-D) Examining the role of JAK/STAT signaling by treating with either 0.1% DMSO (A) or 10µM Ruxolitinib (B) after MTZ treatment from R0-R72. Mi(TnI>GFP) electroporated with Mesp>NTR::mScarlet-I stained for PH3 to label mitotic cells. C) Quantification of the number of PH3$^+$TnI>GFP$^+$ cells per heart in R72 MTZ +0.1% DMSO(R0-R72) versus R72 MTZ +10uM Ruxolitinib(R0-R72). D) Quantification of living/regenerating heart area (TnI>GFP area minus NTR::mScarlet-I debris area) in R72 MTZ +0.1% DMSO(R0-R72) versus R72 MTZ +10uM Ruxolitinib(R0-R72). E-H) Examining the role of Hedgehog signaling by treating with either 0.1% DMSO (E) or 10µM Vismodigeb (F) after MTZ treatment from R0-R72. Mi(TnI>GFP) electroporated with Mesp>NTR::mScarlet-I stained for PH3 to label mitotic cells. G) Quantification of the number of PH3$^+$TnI>GFP$^+$ cells per heart in R72 MTZ +0.1% DMSO(R0-R72) versus R72 MTZ +10uM Vismodigeb(R0-R72). H) Quantification of living/regenerating heart area (TnI>GFP area minus NTR::mScarlet-I debris area) in R72 MTZ +0.1% DMSO(R0-R72) versus R72 MTZ +10uM Vismodigeb(R0-R72). I-L) Examining the role of Hippo/YAP signaling by treating with either 0.1% DMSO (I) or 250nM Verteporphin (J) after MTZ treatment from R0-R72. Electroporated with Mesp>NTR::mScarlet-I stained for MF20 to label myocardium and PH3 to label mitotic cells. K) Quantification of the number of PH3$^+$MF20$^+$ cells per heart in R72 MTZ +0.1% DMSO(R0-R72) versus R72 MTZ +250nM Verteporphin(R0-R72). L) Quantification of MF20$^+$ heart area in R72 MTZ +0.1% DMSO(R0-R72) versus R72 MTZ +10uM Verteporphin(R0-R72). Error bars SEM. * $p<0.05$, ** $p<0.01$, n.s.=not significant. Scale bar: 25µm
